# Mutations of short tandem repeats explain abundant trait heritability in Arabidopsis

**DOI:** 10.1101/2025.03.07.641981

**Authors:** Zhi-Qin Zhang, Juan Jiang, Yong-Chao Xu, Craig Dent, Sridevi Sureshkumar, Sureshkumar Balasubramanian, Ya-Long Guo

## Abstract

Short tandem repeats (STRs) mutations are one of the most important fuels of evolution, but are often largely ignored despite their huge effects on phenotypic variations. Here, we leveraged seven Arabidopsis mutation accumulation lines to assess the STR mutation rate, which ranged from 1.93×10^−2^ to 4.40×10^−3^ per STR locus per generation and was much higher than the common SNP mutation rate at the level of 10^−9^-10^−8^. Interspecific comparison revealed rapid STR turnover, with a large proportion (71.8%) only present in Arabidopsis. In addition, intraspecific comparison of the ten assembled Arabidopsis genomes revealed that 29.3% of STRs were presence/absence variation (PAV), 36.5% of STRs were length variation (LV), 13.0% of STRs were no variation (NV) and the 21.1% STR loci with both LV and PAV. Based on 413 RNA-seq datasets, we identified that length variation of 3,871 STR loci (eSTRs) was associated with gene expression and 651 STR loci (sSTRs) were associated with splicing variation, and over one thousand eSTRs or sSTRs were linked with published genome-wide association study signals. In addition, a direct association between STR length variation and phenotypic variation uncovered a series of STR-trait associations. More notably, using the expression levels of 24,175 genes, the splice strength values of 12,784 splice sites, and 16 phenotypes of natural Arabidopsis populations, we revealed that the STR heritability is 0.111, 0.143 and 0.101, respectively. Overall, our results not only revealed the evolutionary dynamics of STRs, but highlighted the importance of STR variations as an important contributor to missing heritability in the regulation of complex traits.

## Introduction

Short tandem repeats (STRs), also known as microsatellites or simple sequence repeats, are DNA sequence formed by 1 to 6 base pair (bp) units end by end (Ellegren 2004; Willems et al. 2017). STRs can be classified based on their repeat unit size and unit motif. The birth of STRs could be linked to single nucleotide polymorphisms (SNPs) and transposable elements (TEs), resulting from the creation of short repeats (Messier et al. 1996; McGurk and Barbash 2018). STRs, known for their high variability in genomic sequences, often lead to genetic variations in the number of repeat units instead of altering nucleotides (Ellegren 2004). Owing to their significant variability, mutations in STR repeat unit numbers might impact gene expression and function in diverse ways (Gymrek et al. 2016; Fotsing et al. 2019; Wright and Todd 2023).

Although STR mutations are important structural variants, their repetitive nature and short reads from next-generation sequencing have limited their identification and study in-depth (Treangen and Salzberg 2011). Nevertheless, with rapid advances in sequencing technology and analysis pipelines, the mystery of STRs is revealed gradually. First, STRs have been identified and characterized in diverse species, including humans (Willems et al. 2014), penaeid shrimp (Yuan et al. 2021), Arabidopsis (*Arabidopsis thaliana*) (Cao et al. 2014; Press et al. 2018; Reinar et al. 2021; Reinar et al. 2023), maize (Zhao et al. 2023) and *Caenorhabditis elegans* (Zhang et al. 2022). Second, it has been revealed that fluctuations in unit number enable STRs to affect gene function via diverse mechanisms, including modulating epigenetic modification, transcription, and alternative splicing (Conlon et al. 2016; Gymrek et al. 2016; Quilez et al. 2016; Eimer et al. 2018). For example, in Arabidopsis, intronic GAA/TTC triplet expansion within the *IIL1* gene can reduce its expression by inducing 24-nt short interfering RNAs (siRNAs) and repressive histone marks through the RNA-directed DNA methylation pathway (Sureshkumar et al. 2009; Eimer et al. 2018; Sureshkumar et al. 2024). The expansion of this STR leads to an environment-dependent decrease in *IIL1* expression, which substantially hampers this strain’s development (Sureshkumar et al. 2009). In humans, STRs have been demonstrated to be a key regulator of gene expression (Gymrek et al. 2016), and are correlated to a variety of complex human diseases (Verkerk et al. 1991; Conlon et al. 2016; Trost et al. 2020; Malik et al. 2021; Mitra et al. 2021).

Nevertheless, the majority of genome-wide association studies (GWAS) only employ common SNPs for trait association. This has sparked a discussion about the sources of the “missing heritability” of complex traits (Eichler et al. 2010). There are a few factors that could affect the “missing heritability”. First, traditional linkage analysis could not discover rare variations because of their small impact (Fournier et al. 2019). Second, structural variations (SVs) were incompletely detected, leading to an estimation bias caused by incomplete linkage disequilibrium (LD) between genetic markers and causal variants (Mace et al. 2017). Third, epigenetic inheritance in addition to Mendelian heredity of DNA sequences could explain “missing heritability” to some extent (Trerotola et al. 2015). Fourth, complex traits could be affected by environmental factors, making it difficult to accurately measure environmental exposure. Rigorous measurement of the environment and analysis of GWAS data can explain partial missing heritability (Haiman et al. 2007).

A considerable portion of the “missing heritability” may be recovered by introducing STR variation since STRs, as major genetic variations, have a significantly greater polymorphism level and a lower LD with common SNPs (Hannan 2018; Gymrek and Goren 2021). It has been revealed that variable numbers of tandem repeats (VNTRs) are associated with a diverse array of human traits and are casual to several diseases at the copy number level or sequence level (Mukamel et al. 2021). In human, using 44 blood cell and biomarker traits, STRs could explain 5.2-9.7% of GWAS signals (Margoliash et al. 2023), and another study showed that eSTRs (STRs associated gene expression variation) explained 10–15% of the *cis* heritability mediated by all common variants on expression quantitative trait loci (Gymrek et al. 2016). In *C*. *elegans*, eSTRs could increase heritability estimation in 72% of the studied expression traits (Zhang and Andersen 2023). However, STRs have been largely ignored in most association studies.

Arabidopsis has more than 1,000 worldwide re-sequenced genomes, abundant transcriptome and phenotypic data (Kawakatsu et al. 2016; Tabib et al. 2016; The 1001 Genomes Consortium 2016; Durvasula et al. 2017; Zou et al. 2017; Togninalli et al. 2020; Jiang et al. 2024), which is a great model system to study natural variation of STRs (Weigel and Nordborg 2015). Here we leveraged the re-sequenced genomes of seven mutation accumulation lines based on short reads, ten chromosome-level genome assemblies based on long reads, and 1,168 re-sequenced genomes based on short reads, revealed the mutational landscape and the functional impact of STRs in Arabidopsis. In particular, we highlighted the extent to which STR mutations in Arabidopsis could explain the “missing heritability” at gene expression, splicing, and phenotypic level.

## Results

### The mutational landscape of STRs in mutation accumulation (MA) lines

Mutation rate is a general parameter for any genetic element. The mutation rate of nearly every genetic element in Arabidopsis, has been estimated, ranging from SNPs (around 7.00×10^−9^ per site per generation) (Ossowski et al. 2010; Weng et al. 2019) and short indels (1.30×10^−9^ per site per generation) (Weng et al. 2019) to spontaneous epimutations (2.56×10^−4^ per site per generation) (van der Graaf et al. 2015). Polymerase slippage during DNA replication was thought to be the primary source of length variation of STRs (Ellegren 2004). Hereafter, STR mutation rate is the percentage of STR loci that changed their number of repeat units compared to the number of founder accession per generation. However, only the mutation rate of dinucleotide STRs (8.87×10^−4^ per STR locus per generation) has been estimated in Arabidopsis, based on a small number of loci (Marriage et al. 2009). In contrast, in humans and *C. elegans*, STR mutation rates are 5.24×10^−5^ (Steely et al. 2022) and 7.50×10^−5^ (Zhang et al. 2022) per STR locus per generation, which are three and four orders of magnitude higher than those of base substitution of 3.0×10^−8^ (Xue et al. 2009) and 2.7×10^−9^ per site per generation (Denvera et al. 2009), respectively.

To estimate the mutation rates and analyze the characteristics of the mutation of STRs, we leveraged seven published MA line datasets (7-25 generations) derived from seven founders of Arabidopsis (Table S1) (Weng et al. 2019; Weng et al. 2021). STR mutation rates in Arabidopsis range from 1.93×10^−2^ to 4.40×10^−3^ per STR locus per generation (Figure 1A), which is six to seven orders of magnitude higher than SNPs (6.95×10^−9^) and short indels (1.30×10^−9^) (Weng et al. 2019), and much higher than STRs in humans or *C. elegans* (Steely et al. 2022; Zhang et al. 2022). It’s interesting to note that the STR mutation rate varies amongst various MA lines (Figure 1A). This variance may be due to chance or the founders’ genetic background, specifically that the genetic variation between MA lines could involve different pathways correlated with mutation accumulation.

**Figure 1.**
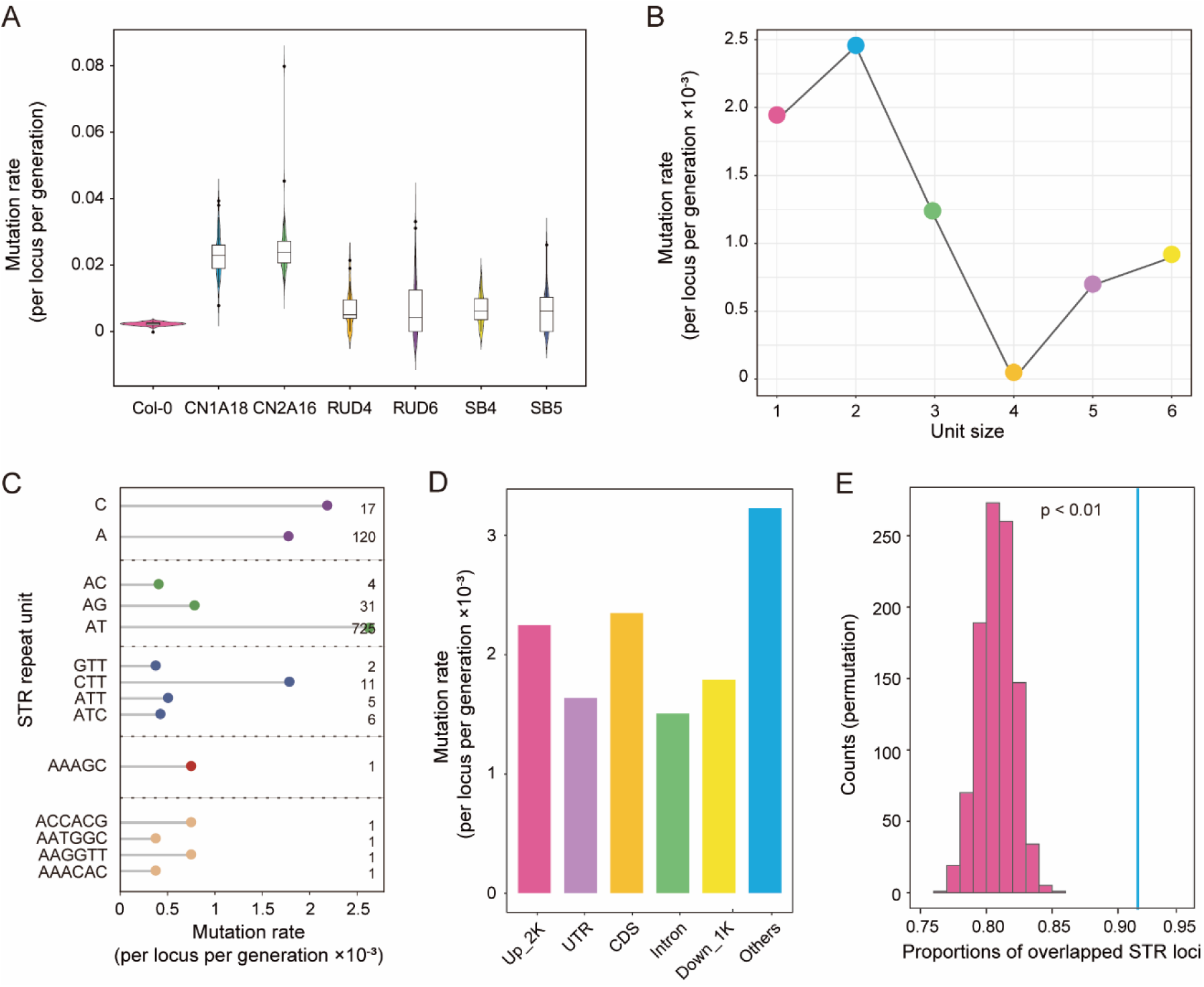
STR mutational landscape in mutation accumulation (MA) lines. (A) STR mutation rates (per STR locus per generation) for seven MA lines. (B) The mutation rates for STRs of different unit size. (C) The mutation rates for STRs of different repeat unit. Some repeat units are not shown because no variation of these motif have been identified in MA lines, probably due to the limited generation. The number on the right represents the number of STR loci identified for different repeat unit. (D) STR mutation rates in different genomic regions. “Others” are STRs not present in 2 kb upstream (Up_2K), untranslated regions (UTR), coding sequences (CDS), intron and 1kb downstream (Down_1K) of genes. (E) The numbers represent the expected and observed overlap ratios between the mutant STR loci found in Col-0 MA lines and the polymorphic STR loci found in the 1,168 natural accessions. The histogram represents the distribution of expected overlap, and 1,000 permutations were performed to generate the expected distribution; and the vertical line represents observed overlap.

To explore the mutation rate variation of different STRs, we used the MA lines derived from the Col-0 accession, which has a larger sample size (107 lines) and propagated more generations (25 generations). After 25 generations, we observed 930 mutations in STRs, and 1,694 single nucleotide mutations. Regarding STRs varying in unit size, dinucleotides exhibited the greatest mutation rate (2.45×10^−3^ per locus per generation), one order of magnitude greater than earlier estimates (8.87×10^−4^ per locus per generation) (Marriage et al. 2009). The previous study estimated based on 54 dinucleotide STR loci in 96 MA lines, whereas our estimation was based on 720 dinucleotide STR loci in 107 MA lines. In contrast, tetranucleotides had the lowest mutation rates (0) among STRs of different unit sizes (Figure 1B). In addition, STRs of different motifs have varied mutation rates, the top motifs with high mutation rate are: C, AT, CTT, AAAGC and AAGGTT (Figure 1C, Table S2). For STRs in different regions of the genome, non-coding regions have the highest mutation rate (3.18×10^−3^ per locus per generation) (Figure 1D), which was also found in single nucleotide mutations (Weng et al. 2019), likely because non-coding regions are under less functional constraints (Monroe et al. 2022). Collectively, these findings imply that variations in STR mutation rates, were influenced by the size of repeat units, repeat motifs, and their genomic location.

To explore whether the STR mutations in MA lines occurred more frequently at polymorphic STR loci (STR loci with length variation) in natural populations, we calculated the observed and expected proportion of MA mutation loci that coincided with the polymorphic STR loci (Figure S1). Among the 930 STR mutation loci identified in the MA lines, 851 (91.5%) overlapped with polymorphic STR loci in natural populations, much higher than the expected ratio based on a random distribution of mutations and polymorphisms (Figure 1E, p < 0.01), suggesting that STR mutations are not random, arise at polymorphic loci more frequently than expected, and there are mutation hotspots of STRs in the genome.

### The landscape of STR mutations in natural populations

To reveal the STR mutation landscape in natural Arabidopsis populations, we utilized ten geographically representative assembled genomes and 1,168 accessions with resequenced genomes (Tables S3 and S4), of which 55 were sequenced in this study (Figure 2A, Table S5), and used its relative species *Arabidopsis lyrata* as outgroup, analyzed the STRs of Arabidopsis at both interspecific and intraspecific levels (Figure 2B). Given long-read sequencing could acquire entire STR alleles (Tanudisastro et al. 2024), we firstly used ten assembled Arabidopsis genomes to perform intraspecific comparisons. Secondly, we used *A. lyrata* as outgroup to perform a comparative study between the two species. Thirdly, we used 1,168 resequenced accessions to characterize the length variation of STRs within the species.

**Figure 2.**
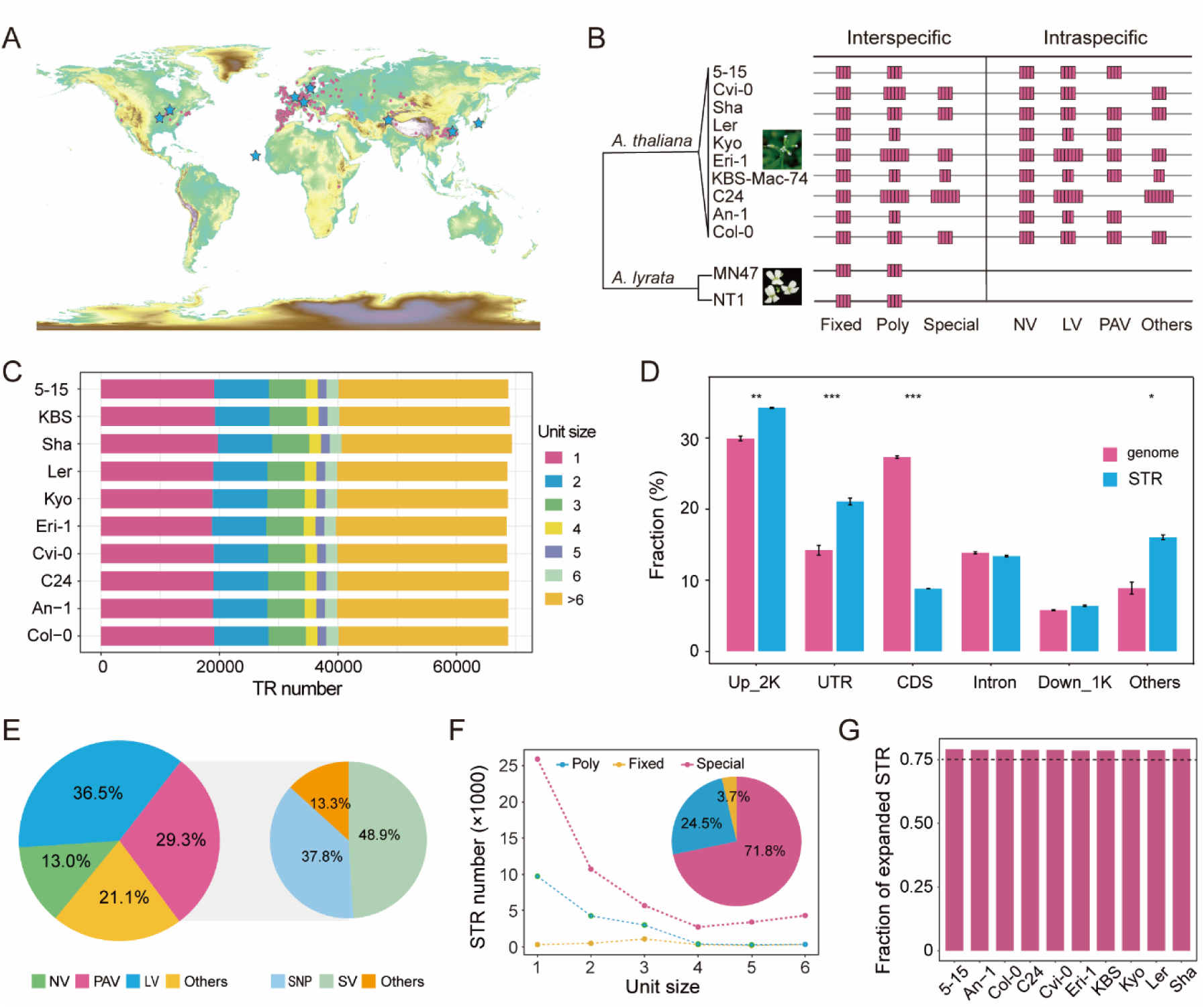
Intra- and interspecific STR mutational landscape. (A) Geographical distribution of nine assembled Arabidopsis non-reference accessions (blue stars) and 1,168 resequenced (red points) accessions analyzed. (B) Schematic diagrams of different classes of STR variation. STR variation among species can be divided into three categories: Fixed, STR loci with the same motif and repeat unit number; Poly, STR loci with the same motif but different repeat unit number; Special, STR loci only present in Arabidopsis. STR variation within species can be divided into four categories: PAV, STR loci with presence/absence variation; LV, STR loci with length variation; NV, STR loci with no variation; Others, STR loci except LV, PAV and NV. (C) Identification of tandem repeats in ten assembled genomes. Each bar shows the relative fraction of homopolymer, dinucleotide, trinucleotide, tetranucleotide, pentanucleotide, hexanucleotide repeats and repeating sequences with repeating units greater than six. (D) Distribution of genomic regions and STR classes across genomic annotations. Genome: the proportion of different functional regions in the ten assembled genomes. STR: the proportion of STR loci in different genomic regions of the ten assembled genomes. Fisher’s exact test was used. *, p < 0.05; **, p < 0.01; ***, p < 0.001. Error bars are means (± s.d.). (E) Left: Overview of the type of STR mutations detected at ten assembled genomes. Right: the proportion of STR PAVs contributed by different variations. (F) Overview of the STR mutation types detected between Arabidopsis and *A. lyrata*. The number of Fixed, Same, and Special STRs for various motif sizes is displayed in a line chart, and the proportions of each are displayed in a pie chart. (G) The proportion of expanded STRs in Arabidopsis relative to *A. lyrata*.

To perform intraspecific comparisons, we first identified STR loci across the Arabidopsis genome by analyzing the Col-0 genome and nine non-reference genome assemblies generated from long-read sequencing by previous studies (Jiao and Schneeberger 2020; Jiang et al. 2024). In the Col-0 reference genome (TAIR10), there are 95,316 tandem repeats identified using Tandem Repeat Finder (TRF v4.09) (Benson 1999), of which 53.8% (51,288 loci) are STRs (unit size is 1-6 bp), comprising 1.5% of the Arabidopsis genome (Figure 2C). In total, 73,717 STR loci were identified across the ten assembled genomes.

Across the ten assembled genomes, STRs accounted for more than half of the tandem repeats; homopolymers were the most abundant type (Figure 2C), similar to those of primates (Verbiest et al. 2022). Among the different genomic regions, STRs were mainly enriched in the regions of untranslated regions (UTR) and 2 kb upstream of genes (Figure 2D). Among different genic regions, the proportion of STRs in coding sequences (CDS) was much lower (Figure 2D), most probably due to the functional constraint in CDS. STR variation in ten assembled genomes of Arabidopsis was classified into four classes: PAV, STR loci with presence/absence variation; LV, STR loci with length variation; NV, no length variation and PAV, and Others, STR loci with both LV and PAV together (Figure 2B). For example, when the length variation of a STR is detected in some accessions, but this STR is not present in other accessions, we define such a STR loci as Others. The proportion of PAV, LV, NV, and Others were 29.3%, 36.5%, 13.0%, and 21.1%, respectively (Figure 2E). There are two main types of mechanisms that directly contribute to PAV: SNPs introduce or disrupt the continuity of repeated sequences; structural variations (SVs) introduce new STRs or remove STRs. In total, 37.8% of PAV can be explained by SNP, and 48.9% of PAV can be explained by SVs. Given the high percentage of PAV, PAV probably could play a more important role than LV.

For interspecific comparisons, we identified STR loci across two genomes of *A. lyrata* (MN47 and NT1) (Wlodzimierz et al. 2023). The comparison of STRs between *A. thaliana* and *A. lyrata* divided STRs of *A. thaliana* into three categories: Fixed, STR loci with the same motif and repeat unit number; Poly, STR loci with the same motif but different repeat unit number; Special, STR loci only present in *A. thaliana* at the syntenic region (Figure 2B). On average, there were 2,731 fixed STRs, 18,082 poly STRs, and 52,904 special STRs in *A. thaliana*. Apparently, Special STRs was the most abundant (>70.0%), indicating that STRs gained or lost rapidly between species (Figure 2F). Interestingly, within the special STRs, homopolymers occupied nearly more than half of all the special STRs, reaching 25,964 (Figure 2F).

Given that STR mutations caused by DNA slippage are more likely to cause length variation, it is interesting to know whether most poly STRs expanded or contracted in *A. thaliana* compared to its sister species. To address this question, we counted the proportion of expanded and contracted STRs in Arabidopsis relative to the *A. lyrata* reference genome (MN47), based on ten assembled genomes of *A. thaliana*. More than 75.0% loci are expanded in all ten accessions in comparison to *A. lyrata* reference genome (MN47) (Figure 2G), which is consistent with the reduction in the effective population size of the *A. thaliana,* making the expanded STRs more likely to become fixed (Hu et al. 2011). Taken together, there were more expanded STRs than contracted STRs in *A. thaliana* compared to its sister species *A. lyrata*.

To characterize the STR mutations in natural populations, we used HipSTR (Willems et al. 2017) to identify the length variation of STRs in 1,168 resequenced Arabidopsis genomes. Of the 51,288 STR loci in the reference genome, 40,824 STR loci (79.6%) have length variation across the 1,168 accessions. Using four accessions (5-15, Ler, Cvi-0, Kyo) with both long-read genome assemblies and resequencing reads, we performed a validation analysis for STRs identified in resequenced genomes, and found that over 96% of the STRs were consistent in all the four accessions (Table S6). This high consistency indicates a high accuracy of variant calling of STRs in resequenced genomes.

Among STRs of different repeat unit length, homopolymers were the most polymorphic STRs (pSTRs) (Figure S2). For all pSTRs together, each STR locus has on average 4.2 alleles (Figure 3A), similar to a previous study (4.5 alleles) (Press et al. 2018). STRs have a pivotal role in contributing to the diversity within and between populations (Xie 2024). Here, among nine non-relict populations, only a small fraction (1.3%) of pSTR loci have length variation across all populations, suggesting a high intraspecific turnover rate of STRs (Figure 3B). Furthermore, we discovered that Yangtze River basin population has the most abundant unique STRs, which could result from a higher STR mutation rate or reduced purifying selection accompanying the expansion to this new habitat (Figure 3B).

**Figure 3.**
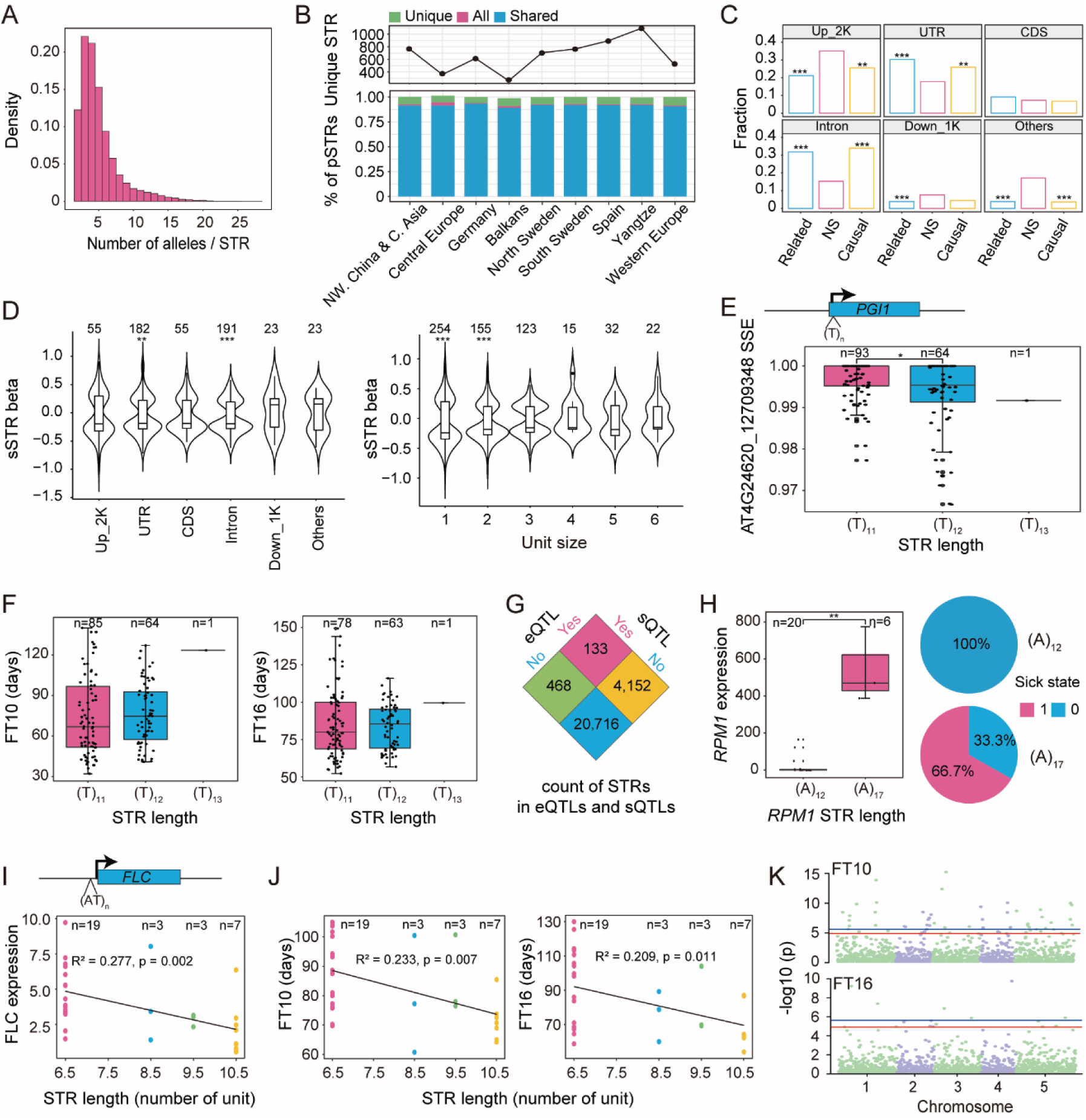
The functional impact of STR variation. (A) The distribution of allele counts per STR locus across all genotyped STRs in 1,168 accessions. (B) Number of unique STRs (upper) and proportions of different polymorphic STRs (pSTRs) (lower) across different populations of 1,168 accessions. Unique, only exist in one population; Shared, exist in two or more but not all populations; All, exist in all populations. (C) The proportion of sSTRs in each category (NS, related, or causal) in each genomic region. NS, STRs that were not reported to be significantly correlated with any alternative splicing events; related, STRs that were significantly associated with at least one alternative splicing event but were not reported as causal; causal, STRs that were reported as causal for at least one alternative splicing event. The p value displayed above the bars represents Fisher’s exact test for enrichment significance, in contrast to STRs, which showed no significant correlation with other splicing events. *, p < 0.05; **, p < 0.01; ***, p < 0.001. (D) Left: the effect size (beta value) of sSTRs classes across genomic annotations. Right: the effect size of sSTRs classes with different motif lengths. Binomial test was used. Beta value represents the effect size of STR loci on splicing variation, and a value greater than 0 indicates a positive correlation. (E) sSTR associated with AT4G24620_12709348 splice variation. The x-axis shows STR genotype and the y-axis gives Splice-site Strength Estimate (SSE) value. (F) Repeat number variation of STR in the intron of *PGI1* is correlated with flowering time variation. The x-axis shows STR genotype and the y-axis gives flowering time in 10°C (FT10) and 16°C (FT16). (G) Count of STR in eQTL and sQTL. eQTL: expression quantitative trait loci. sQTL: splicing quantitative trait loci. (H) *RPM1* repeat number variation is linked with GWAS signal of pathogen susceptibility. Left: the x-axis shows STR genotype and the y-axis gives normalized *RPM1* expression. Wilcoxon test was used. Right: the pie chart represents the proportion of different susceptibility states under (A)_12_ (upper) and (A)_17_ (lower). 0 represents disease resistance and 1 represents disease susceptibility. (I) eSTR associated with *FLC* expression variation. The x-axis shows STR genotype and the y-axis gives normalized *FLC* expression. (J) Repeat number variation of STR in 2 kb upstream of *FLC* is correlated with flowering time variation. The x-axis shows STR genotype and the y-axis gives flowering time in 10°C (FT10) and 16°C (FT16). (K) GWAS analysis of FT10 and FT16. The red horizontal line corresponds to the significance threshold (0.05/STR loci number) and blue horizontal line corresponds to the significance threshold (0.01/STR loci number).

### The impact of pSTR length variation on gene expression and alternative splicing

To investigate the impact of STR length variation on gene function, we focused on 413 accessions with transcriptome data. A previous study has demonstrated that STR length variation could affect gene expression (Reinar et al. 2021). We first identified STR loci whose length variation was associated with adjacent gene expression variation (eSTRs). Intriguingly, of these 3,871 eSTRs, 58.8% eSTRs mainly upregulated the expression of nearby genes with the elongation of STR length (Figure S3A), regardless of their location. In terms of the repeat unit sizes, except for tetranucleotides and hexanucleotides, other unit sizes are more inclined to elevate the expression of genes with the increased STR length (Figure S3B). Given that the eSTRs identified above could potentially be explained by their tagging with nearby other variants (SNPs/indel/TEs), to prioritize potentially causal eSTRs, we further employed CAVIAR (Hormozdiari et al. 2014) to quantify the posterior probability of causality for each variant. A total of 15.9% of eSTRs (615) with the highest posterior probability of causality among all the adjacent variants tested were regarded as potential causal eSTRs (FM-eSTRs). Here all STRs were grouped into three categories: not an eSTR for any gene (NS), identified as an eSTR but not a FM-eSTR (related), and identified as a FM-eSTR (potentially causal). To explore the potential effect of STR location on gene expression, we compared eSTR (causal and related) to the non-eSTR set (NS). eSTRs were enriched in UTR, CDS and intron of the associated genes (Figure S3C). Genes with expression level variation associated with STRs (eGenes) were significantly enriched in the stimulus response pathway, regulation of biological, and other pathways (Figure S3D). Given the stimulus response is largely associated with the adaptation to climate change in a short period of time, STRs is one of the rapidly evolved genetic elements in genome, which may be the reason that why eGenes is significantly enriched in stimulus response pathway.

STRs can affect the alternative splicing (AS) of genes (Press et al. 2018; Sulovari et al. 2019; Hamanaka et al. 2023), therefore, we estimated the effect of STRs on splicing by examining their effects on the strength of each splice site (Dent et al. 2021). Splice site strength was calculated among RNA-seq reads mapping to each splice site, as the proportion of reads which provide evidence for the usage of that splice site, using SpliSER v0.1.8 (Dent et al. 2021). In total, we identified 354 splicing STRs (sSTRs), whose length variation was associated with the splice site strength of 601 distinct splice sites from 295 genes with FDR < 10%. Consistent with eSTRs, sSTRs that could be interpreted by their tagging with nearby variants (SNPs/indels/TEs) were filtered out, and 195 sSTRs were identified as potentially causal sSTRs (FM-sSTRs), accounting for 55.1% of the total sSTRs.

To explore the potential effects of STR location on splice site strength, we examined the enrichment of sSTRs relative to functional genomic elements. We grouped STRs into three categories: not an sSTR for any gene (NS), identified as an sSTR but not a FM-sSTR (related), and identified as a FM-sSTR (potentially causal). Compared to the NS set, sSTRs (both related and potentially causal) were enriched in UTR and intron of associated genes (Figure 3C), as expected that STRs in UTR and intron have a strong impact on the alternative splicing. Regardless of their unit size, these 354 sSTRs are more likely to strengthen splice sites as they elongated (Figure 3D). In terms of the location of STRs, STRs in upstream region and gene bodies were tend to promote alternative splicing with increased length (Figure 3D). Intriguingly, we found an FM-sSTR fell in the intron of *PGI1*(AT4G24620), a positive regulator of flowering time; and its length was positively correlated (p < 0.05) with the splicing efficiency of AT4G24620 and flowering time (Figure 3E and 3F).

Changes in splicing can potentially lead to nonsense-mediated decay (NMD), which can affect gene expression level. Therefore, we identified STRs that can affect splicing and gene expression simultaneously. There were 133 STRs were linked to the same gene’s AS and expression (Figure 3G). In particular, we found a fine-mapped sSTR within intron of *IIL1*, whose length was negatively correlated with *IIL1* expression and showed a positive relationship with splice strength of AT4G13430, and this STR has been demonstrated to have functional effects (Sureshkumar et al. 2009), providing supporting evidence for our identification accuracy. Overall, these results indicated that the impact of STRs on alternative splicing is correlated with their expression levels.

### The contribution of STR length variation to phenotypic variation

To assess the impact of STR length variation on phenotypic variation, we measured it in two dimensions. In human, it has been revealed that STRs could explain 5.2-9.7% of GWAS signals (Margoliash et al. 2023), and a previous study suggested that the causal loci of GWAS signal are STR mutations (Grünewald et al. 2015). Therefore, firstly, we leveraged the published GWAS catalog (Togninalli et al. 2020) to identify STR loci linked with previously identified GWAS signals (r^2^ > 0.2). Secondly, we performed direct association analysis between STR length and phenotypes.

In total, we identified 9,629 STR loci linked with 13,004 GWAS signals, 1,500 eSTRs linked with 8,334 GWAS signals, and 354 sSTRs linked with 3,193 GWAS signals. For STR loci linked with GWAS signal, a noncoding STRs lies in the 1 kb downstream of the *RPM1*, a gene controlling the susceptibility of Arabidopsis to *Pseudomonas syringae*, and is strongly linked with the published GWAS signal of *P. syringae* resistance (chr3: 2229952, r^2^ = 0.75). The accessions with longer STR (A_17_) have higher expression level of *RPM1* than those with shorter STR (A_12_) (Figure 3H). The published phenotypic data (Atwell et al. 2010) suggested that Arabidopsis with longer STR (A_17_) at the 1 kb downstream of the *RPM1* are probably more susceptible to *P. syringae* than those with shorter STR (A_12_) (Figure 3H).

For eSTRs co-localized with published GWAS signal, an FM-eSTR falling in 2 kb upstream of *FLC* was linked with the GWAS signal (chr5: 3183501, r^2^ = 0.39) of flowering time at both 10°C and 16°C, and the length of eSTR was negatively correlated with *FLC* expression (Figure 3I). For the association between this FM-eSTR polymorphisms and flowering time, we found that the longer eSTR length is associated with earlier flowering at both 10°C and 16°C conditions (Figure 3J).

Given STRs weakly linked with GWAS signal could associated with phenotypic variation, we used associaTR (Mousavi et al. 2020) to directly identify STRs that are associated with phenotypes. Using 16 phenotypes which have been measured in over 200 accessions (Table S7), we identified 12 and 54 significant STRs that are associated with flowering time variation but not co-localized with flowering time GWAS signal at 16°C and 10°C, respectively (Figure 3K). Numerous noteworthy STRs were also linked to other traits, indicating their influence on the phenotypes under study (Figure S4).

### The contribution of STR length variation to trait heritability

STR variations were considered to have an essential role in the heritability of complex traits in humans and other model species (Mukamel et al. 2021; Zhang and Andersen 2023). However, most GWAS analyses did not include STR variation, which might lead to the missing heritability to some extent (Phadte et al. 2023).

To test the potential of STR length variation in explaining missing heritability, we compared the LD of SNP-SNP and STR-SNP, respectively. It turns out that the LD of STR-SNP was much lower than that of SNP-SNP, indicating that traditional GWAS signal could not tag most STRs (Figure 4A).

**Figure 4.**
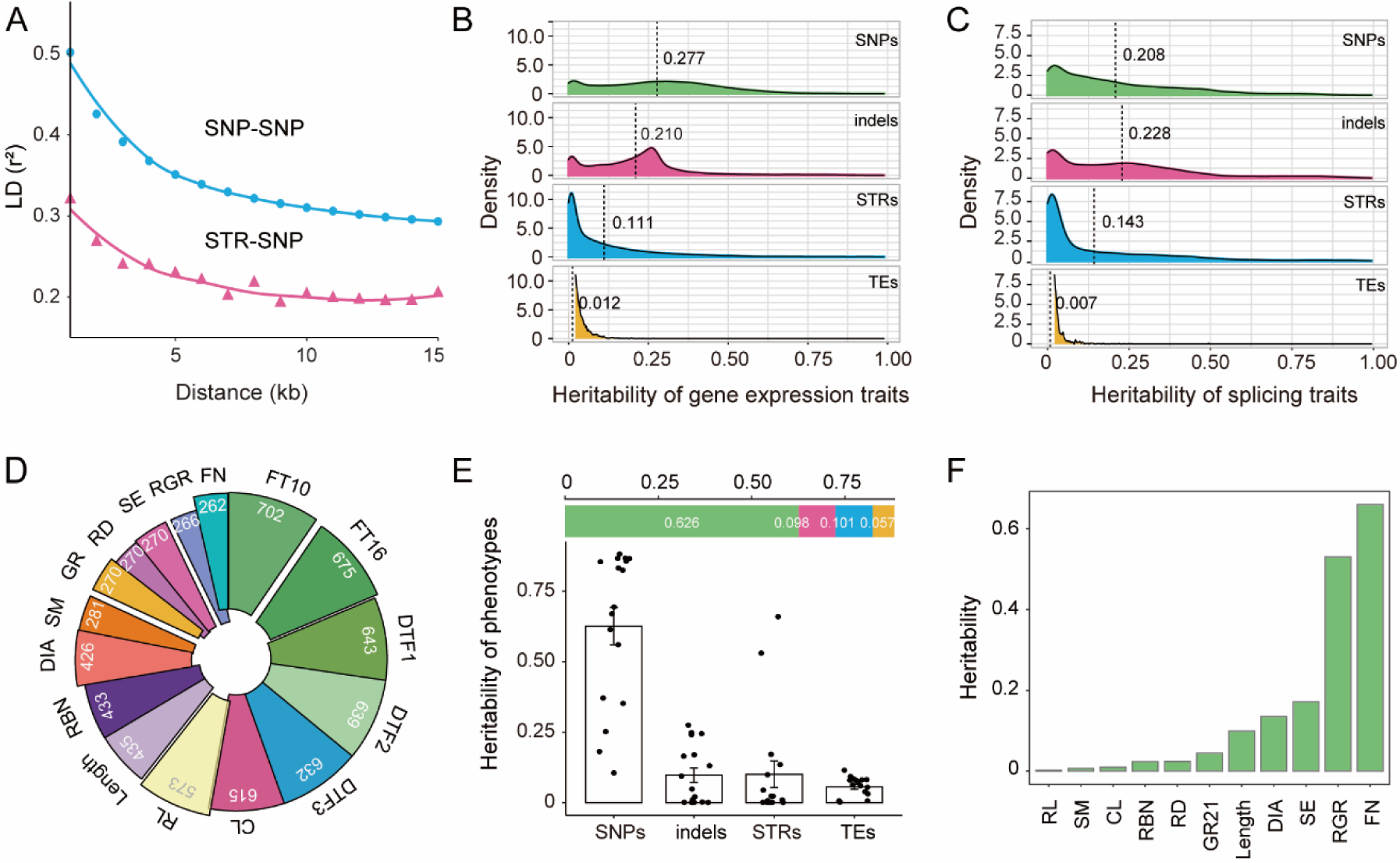
Trait heritability contributed by STRs. (A) LD estimates for SNP-SNP and SNP-STR loci. Loess regression lines for each category are plotted. (B) Comparison of heritability of gene expression level contributed by STRs, SNPs, TE and indels. Heritability was estimated by random effects corresponding to each category. The vertical dashed lines indicate the mean values. (C) Comparison of heritability of alternative splicing efficiency contributed by STRs, SNPs, TEs and indels. The vertical dashed lines indicate the mean values. (D) The phenotypes used to calculate heritability. Numbers indicates available accessions of each phenotype. (E) Comparison of heritability of phenotypes contributed by STRs, SNPs, TEs and indels. Horizontal bar chart represents the average heritability contributed by various variants; from left to right, SNPs, indels, STRs and TEs. (F) The heritability contributed by STRs in diverse phenotypes. Four phenotypes (DTF2, CL, RL and DIA) without any heritability contribution from STRs were not shown.

To estimate the contribution of STRs to missing heritability, we used two strategies. Firstly, using the LDAK-thin model, we compared heritability explained with and without STRs model to determine whether STRs could capture missing heritability. Secondly, we partitioned the relative contributions of STRs in comparison to those of SNPs, indels, and TEs to determine the contribution of STRs to heritability (Speed et al. 2020). The latter approach has been utilized in multiple studies to quantify the overall contribution of variations to the heritability of quantitative traits, and to categorize the contributions of different classes of variants (Yang et al. 2010; Gymrek et al. 2016; Zhou et al. 2022). We estimated heritability from three dimensions: gene expression, alternative splicing and phenotype.

Using the first strategy, the average heritability of expression increases from 0.579 to 0.610, and the average heritability of phenotype increases from 0.865 to 0.882, when STRs are included in the model compared with model that consider only SNPs, indels and TEs, indicating that STRs capture partially missing heritability.

To partition the contribution of STRs versus other variant types to heritability, at the gene expression level, we used expression traits of 24,175 genes based on 413 accessions with both high-coverage resequencing and RNA-sequencing data. The composite model estimated that STRs contributed to 11.1% of the phenotypic variance, higher than TEs (1.2%), but lower than SNPs (27.7%) and indels (21.0%) (Figure 4B). At the alternative splicing level, we used splicing traits of 12,784 splice sites based on 413 accessions and found that 14.3% of the phenotypic variance can be explained by STRs, in comparison to SNPs (20.8%), TEs (0.7%) and indels (22.8%) (Figure 4C). At the phenotypic level, we used 16 phenotypes that each have data on over 200 accessions (Figure 4D), and found that STRs contributed to 10.1% of the phenotypic variance, in comparison to SNPs (62.6%), TEs (5.7%) and indels (9.8%) (Figure 4E). Furthermore, in terms of the differential contribution of STRs in diverse phenotypes, we found that STRs could explain a much higher variation in Fruit number (FN), up to 65.97% (Figure 4F). Taken together, heritability in expression variation, splicing variation, and phenotypic variation can be partially explained by STR variants, which must be an important contributor to missing heritability. However, the explanation power differed among traits, most probably depending on their genetic architecture.

## Discussion

Short tandem repeats (STRs) represent one of the main sources of genetic diversity across a wide range of species. In order to investigate how STRs evolved and how length variations could impact gene function and contribute to phenotypic variation, we performed a global analysis of STRs in natural Arabidopsis populations. Although in Arabidopsis, the variation and function of STRs has been studied previously (Press et al. 2018; Reinar et al. 2021). Nevertheless, compared to the prior analysis (McGurk and Barbash 2018), we employed a larger data set here. Specifically, in order to investigate the functional effects of STR length variation, we also addressed its impacts on splicing variations and phenotypes in addition to the expression variation from the previous study (Reinar et al. 2021). Most importantly, in this study, we estimated the contribution of STRs to heritability at different phenotypic context in Arabidopsis.

In general, the repetitive nature of STRs and short reads from next-generation sequencing limited their identification and understanding. Here we integrated the de novo assembled genomes based on long reads sequencing of ten representative natural accessions, and 1,168 resequenced genomes of natural populations and seven mutation accumulation lines based on short reads, and performed inter- and intra-specific comparison to understand the evolutionary dynamics of STRs. In particular, we systematically analyzed STR mutations using seven MA lines and found that STR mutation rates were much higher than the rate of gene gains and losses (0.001359 per gene every million years) (Guo 2013), SNPs (6.95×10^−9^ per site per generation) and short indels (1.30×10^−9^ per site per generation) (Weng et al. 2019) and spontaneous epimutations (2.56×10^−4^ per site per generation) (van der Graaf et al. 2015), and were influenced by a variety of factors, such as genetic background, location on the genome, unit motif, and the unit size. The mutation rate of STRs in Arabidopsis is similar to that of *Daphnia magna* (Ho et al. 2019). Although TE mutation rate is not available for MA lines or natural populations in Arabidopsis, but it has been estimated in *Daphnia magna* at the rate of 10^−5^ (per site per generation) (Ho et al. 2021), and in *Drosophila melanogaster* at the rate of 10^−9^ (per site per generation) (Nuzhdin and Mackay 1995; Adrion et al. 2017; Ho et al. 2019). In addition, previous STR variation studies mainly focused on length variation, here we used assembled data and found that PAV is a hidden form of STR variation, accounted for 29.3% of the overall variation, suggesting that utilizing genome assembly of multiple accessions would capture the full landscape of STR mutations in natural populations and facilitate the functional characterization of STR mutations.

Missing heritability is defined as the discrepancy between the variance explained by all significant variants in GWAS and the heritability estimates from familial-based genetic studies (Maher 2008; Manolio et al. 2009; Eichler et al. 2010; Press et al. 2014; Zhou et al. 2022). Missing heritability could be partially due to the missing of STR loci in most GWAS analyses (Gymrek et al. 2016; Hannan 2018; Mukamel et al. 2021). A considerable portion of the “missing heritability” may be recovered by introducing STR variation since STRs, as major genetic variations, have a significantly greater polymorphism level and a lower LD with other genetic markers. For example, variable numbers of tandem repeats (VNTRs) are associated with a range of human traits and diseases, either at the copy number level or sequence level (Mukamel et al. 2021). In human, using 44 blood cell and biomarker traits, STRs were found to drive 5.2-9.7% of GWAS signals for these traits (Margoliash et al. 2023).

To figure out the contribution of STRs to missing heritability, for the first time, we estimated the heritability of STR length variation on gene expression level, splice strength and phenotypes in Arabidopsis at the same time. For the gene expression level, splicing strength and phenotypes, the average STR heritability was 0.111, 0.143 and 0.101, respectively. In terms of contribution of STRs to the expression variation, here in Arabidopsis the contribution of STRs to expression variation is much higher than that in human (11.1% vs 1.8%) (Gymrek et al. 2016). Overall, STRs is the most rapidly evolving genetic components in the genome and must be an important contributor to missing heritability. STRs can quickly produce a lot of genetic variations, which could be crucial for rapid adaption. Much more in-depth analyses are needed to clarify the dynamics of STRs and its roles in evolution.

Nevertheless, a few issues should be considered in further studies. 1) Although we used data from 10 assembled genomes from long-read sequencing in addition to the short-read sequencing data to assess the STR mutation landscape, more samples with long-read sequencing might benefit this type of study. 2) Given limited samples with phenotypic data used here, there is noise when estimating the contribution of STR to phenotypic heritability. 3) Although there is substantial evidence that the candidate casual STRs co-localized with published GWAS signals, rigorous molecular biology confirmation will be crucial. Overall, in-depth study of STRs will be highly beneficial to both understanding of its evolutionary process and its effect on evolution of diverse species.

## Materials and methods

### Plant material and high-throughput DNA sequence

55 Arabidopsis natural accessions sequenced in this study were collected from the Yangtze River basin (Table S5). Genomic DNA was extracted from the seedlings using the CTAB method. Paired-end sequencing libraries were constructed with insert size around 350 bp. Illumina HiSeq X Ten was used to generate the 150 bp paired-end reads.

### STR identification and classification

The reference genome TAIR10 was used to produce a reference set of short tandem repeats (STRs) with the TRF software (v4.09) (Benson 1999) and further filtered as described in a previous study (Fotsing et al. 2019). After filtering, we retained a reference set with 51,288 STRs. Using the same approach, we identified 104,153 and 96,848 STRs from the two *A. lyrata* genomes MN47 and NT1, respectively.

To identify non-reference STRs, we used nine Arabidopsis assemblies and the reference Col-0 (Table S3). First, we performed pairwise whole genome alignment among the ten genomes with AnchorWave (Song et al. 2022). Second, we identified STRs in nine assembled genomes using the same method as the reference genome analysis. Then, STRs that do not exist in the reference genome were integrated together with a reference set to form the Arabidopsis panSTR datasets.

PAV is presence/absence of a STR locus. To understand the causes of STR PAV formation, we classified STR PAV into three categories: SNP, SV and other. PAV is referred to as SNP-induced if SNPs induced its presence/absence, SV induction if SV induced its presence/absence, and other induction if other mutation induced its presence/absence.

In total, 1,168 resequenced natural accessions were used in this study: 810 accessions from 1001 Genomes Project (The 1001 Genomes Consortium 2016), 60 accessions from published data of Africa (Durvasula et al. 2017), 298 accessions from China and sequenced by our laboratory, of which 243 accessions were published previously (Zou et al. 2017; Jiang et al. 2024). All these accessions were sequenced with at least 10x coverage. BWA was first used to align each accession’s reads to the TAIR10 reference genome (Li and Durbin 2009). Next, we used HipSTR to call polymorphic STRs in all 1,168 sequenced accessions (Willems et al. 2017). All samples were genotyped together with the parameters “--min-reads 5 and --def-stutter-model”. VCFs were filtered using the filter_vcf.py script available from HipSTR, using recommended settings (--min-call-qual 0.9, -- max-call-flank-indel 0.15 and --max-call-stutter 0.15). After filtering, we got a matrix which containing 1,168 columns (accessions) and 40,824 rows (STR loci).

To assess the identification accuracy of STR length variation in resequenced genomes, we used four long-read assembled genomes (5-15, Ler, Cvi-0, Kyo) and extracted sequence from the orthologous regions corresponding to Col-0 STRs. Furthermore, we compared these extracted sequences with those identified by the corresponding short read data, and verified them when the sequences were completely consistent. Finally, we counted the number of consistent STR loci identified with both long-read genome assembly and short-read data, and computed the identification accuracy of each accession.

### Identification of STR mutations from accumulation lines

We leveraged seven existing mutation accumulation (MA) line resequencing datasets (7-25 generations) derived from seven founders from natural Arabidopsis populations of French and Swedish (Table S1), including founder CN1A18: 56 lines for 10 generations; CN2A16: 51 lines for 10 generations; RÖD4: 50 lines for 8 generations; RÖD6: 50 lines for 8 generations; SB4: 53 lines for 8 generations; SB5: 56 lines for 8 generations and founder Col-0: 107 lines for 25 generations. The identification method of polymorphic STR was the same as that used for 1,168 resequenced accessions. The STR mutation rate μ was calculated as m/2nt (Denvera et al. 2009) for each MA line, where m is the number of the mutation (STR unit number changed in comparison to the founder accession), n is the total number of STR loci between the two lines, and t is the number of generations.

### Identification of STR loci correlated with gene expression variation

Transcriptome data of 413 accessions were downloaded from NCBI GEO GSE80744 (Kawakatsu et al. 2016). The STRs of these 413 accessions were retrieved from a STR matrix including 1,168 accessions, and all STR loci with minor allele dosages of less than 3 were excluded. A total of 34,969 STR loci remained for expression analysis. Following a previous research (Fotsing et al. 2019), for each STR locus within 2 kb range of a gene, we performed a linear regression between STR lengths and the normalized gene expression values. Then, we used an FDR (false discovery rate) threshold of 10% to identify significant STR-gene pairs. Altogether, we identified 3,871 unique eSTRs associated with the expression variation of 4,285 genes. Expressions of SNPs, indels and TEs (eSNPs, eindels, eTEs) were identified using the same model covariates and normalization procedures, but using biallelic alleles (0 or 2) rather than STR lengths.

### Identification of STR loci correlated with splice variation

We used SpliSER (Dent et al. 2021) to quantify individual splice-site strengths from Arabidopsis accessions. Splice site strength was calculated among RNA-seq reads mapping to each splice site, as the proportion of reads which provide evidence for the usage of that splice site. The splice-strength data was used to analyze the impact of STR polymorphisms on splicing. Similar to the eQTL analysis, for each STR within 2 kb range of a gene, we performed a linear regression between STR lengths and SSE values. The cutoff for significant sQTLs was set to 10% FDR. Alternative splicing-associated SNPs, indels, and TEs were identified using the same way as we performed with STRs.

### Fine-mapping of causal variants for each eGene and splice site

CAVIAR (ver. 2.1) (Hormozdiari et al. 2014) was used to identify potential causal variants in the associated region for each eGene. CAVIAR generates a list of potential causative variants along with their posterior probability based on two inputs: a linkage disequilibrium (LD) file and a z-score file. Pairwise LD between the eSTR and eSNPs/eindels/eTEs were estimated using the Pearson correlation between dosages (0, 2) and STR genotypes across all samples. CAVIAR was run with parameters “-f 1 -c 2” to model up to two independent causal variants per locus.

Expression-associated SNPs, indels, and TEs were identified using the same model covariates and normalization procedures, but using dosages (0, 2) rather than STR length. The SNPs and indels were called with GATK pipelines (GATK v2.1.8) (DePristo et al. 2011) and TEs were obtained from one of our recent study (Jiang et al. 2024). All of these variants with MAF > 5% were used in expression analysis. On average, for each gene, we tested the correlation of gene expression with the variation of 0.9 STRs, 47 SNPs, 8 indels, and 0.5 TEs. To identify potential causal STR associated with splice site strength, similar analyses were performed as the eQTL analysis using CAVIAR.

### Colocalization of STRs with published GWAS signals

Published GWAS signals were obtained from the Arabidopsis GWAS catalog (Togninalli et al. 2020). Linkage disequilibrium (LD) between STRs and SNPs was computed by taking the squared Pearson correlation between STR lengths and SNP dosages for each STR–SNP pair. To ensure sufficient statistical power, we only retain common SNPs (MAF > 5% in 1,168 accessions). If minor allele dosage less than 3, the STR was excluded from LD calculations (Fotsing et al. 2019). STRs with LD value (r^2^) greater than 0.2 were regarded as the GWAS signal-colocalized STRs.

### Genome-wide association analyses

A total of 16 phenotypes with each has over 200 matched accessions were downloaded from AraPheno (Table S7 and Figure 4B) (Togninalli et al. 2020). We normalized each phenotype as previously described (Margoliash et al. 2023). We performed STR association test using associaTR (Mousavi et al. 2020). AssociaTR fit the linear model y = g * β_g_ + C * β_c_ + ɛ, where y is the vector of phenotype values per individual, g is the vector of STR length dosage genotypes per individual. To select loci with minor allele dosage less than 20, we employed the flag “--mac 20.” In the GWAS study, biallelic SNPs with MAF > 5% and missing rate < 10% were employed. The significance cutoff was set to 0.01/total STR loci or 0.05/total STR loci based on Bonferroni correction.

### Heritability calculation

Before calculating the heritability of STRs, we split each multiallelic STR into multiple biallelic loci. For each STR locus, we decomposed it into the corresponding number of loci according to the number of alleles it contained. We redefined the genotype of each sample at each locus. For each allele: if the genotype is the same as the reference, it is redefined as 0, otherwise, it is defined as 1.

LDAK-thin model (Speed et al. 2020) was used to estimate the proportion of phenotypic variance explained by STRs, SNPs, TEs and indels. First, LDAK was run with parameters “--window-prune 0.98 and --window-kb 100” to exclude nearby SNPs in perfect LD. Then, the kinship matrix was calculated with the power parameter “--alpha −0.5” (Zhou et al. 2022), which determines the expected relationship between per-variant heritability and MAF. Finally, we estimated the kinships for different types of genetic variants (STRs, indels, TEs and SNPs) and used a composite model to calculate the heritability. A parameter “--constrain YES” was added to ensure that heritability ranges from 0 to 1. A total of 24,175 gene expression traits, 12,784 splicing traits and 16 phenotypes containing 7,391 phenotypic values were used to estimate the heritability explained by different variant types.

### Statistical analyses

All the statistical analyses were performed with R (http://www.r-project.org/).

## Data accessibility

Raw sequence data can be accessed at the Genome Sequence Archive in National Genomics Data Center, China National Center for Bioinformation (GSA: CRA012396) that are publicly accessible at https://ngdc.cncb.ac.cn/gsa.

## Acknowledgements

We would like to thank Melissa Gymrek (University of California San Diego), Yao Zhou (Institute of Botany, Chinese Academy of Sciences), and Jia-Fu Chen (Southwest Forestry University) for helpful suggestions about the data analysis. Especially, we thank the anonymous reviewers for their valuable comments that improve our manuscript a lot. This work was supported by National Natural Science Foundation of China (31925004 and 32430008) to Y-L.G., Australian Research Council (ARC) Future Fellowship (FT190100403) to SS, and ARC Discovery Project Grant (DP190101479) to SB.

