## Supplement for "Mutations of short tandem repeats explain abundant trait heritability in Arabidopsis"

Running title: STRs explain abundant trait heritability

Key words: *Arabidopsis thaliana*, evolution, missing heritability, mutation rate, natural variation, short tandem repeats

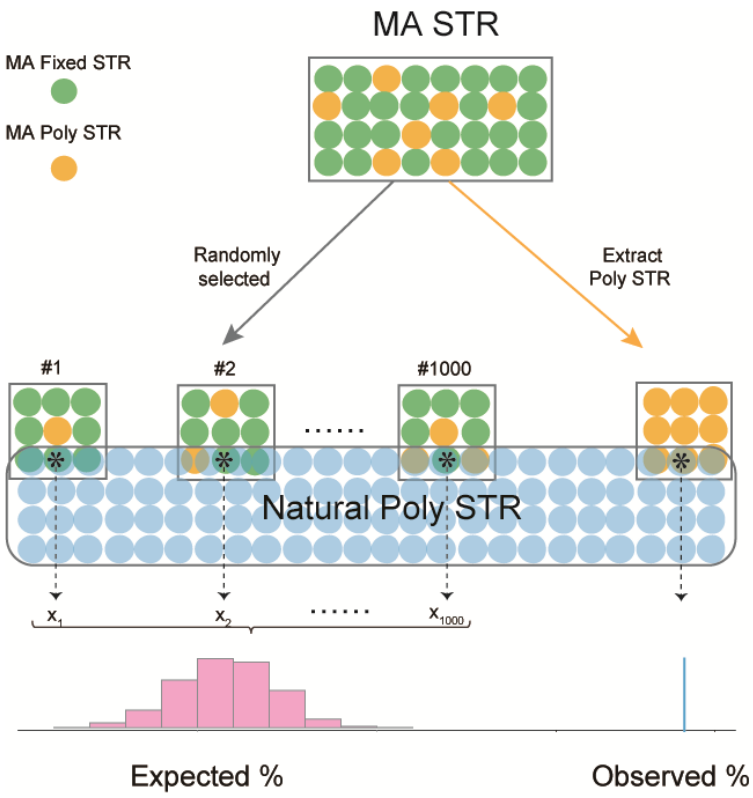


Figure S1. De novo STR mutation preference analysis schematic. Natural Poly STR: STR loci with length variation in 1,168 accessions. MA Fixed STR, STR loci without variation; MA Poly STR, STR loci with length variation in mutation accumulation lines (MA). Observed %: the proportion of MA Poly STR that intersects with Natural Poly STR. Expected %: the proportion of randomly selected MA STR that overlaps with Natural Poly STR.

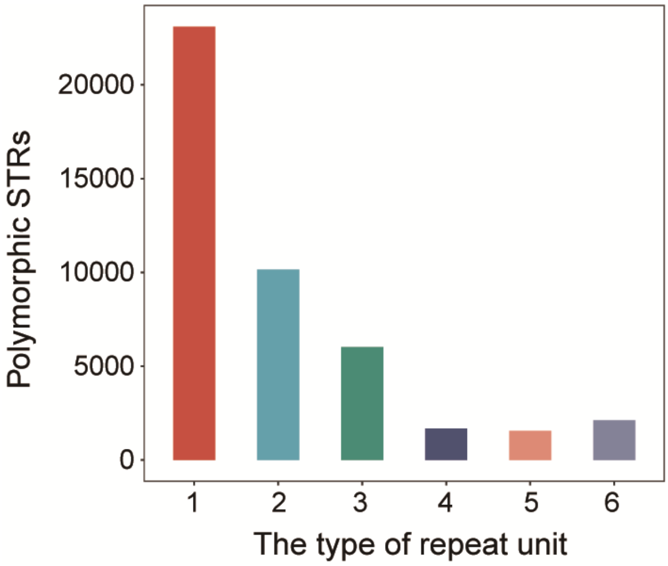


Figure S2. Number of polymorphic STR loci with different unit length identified in 1,168 accessions.

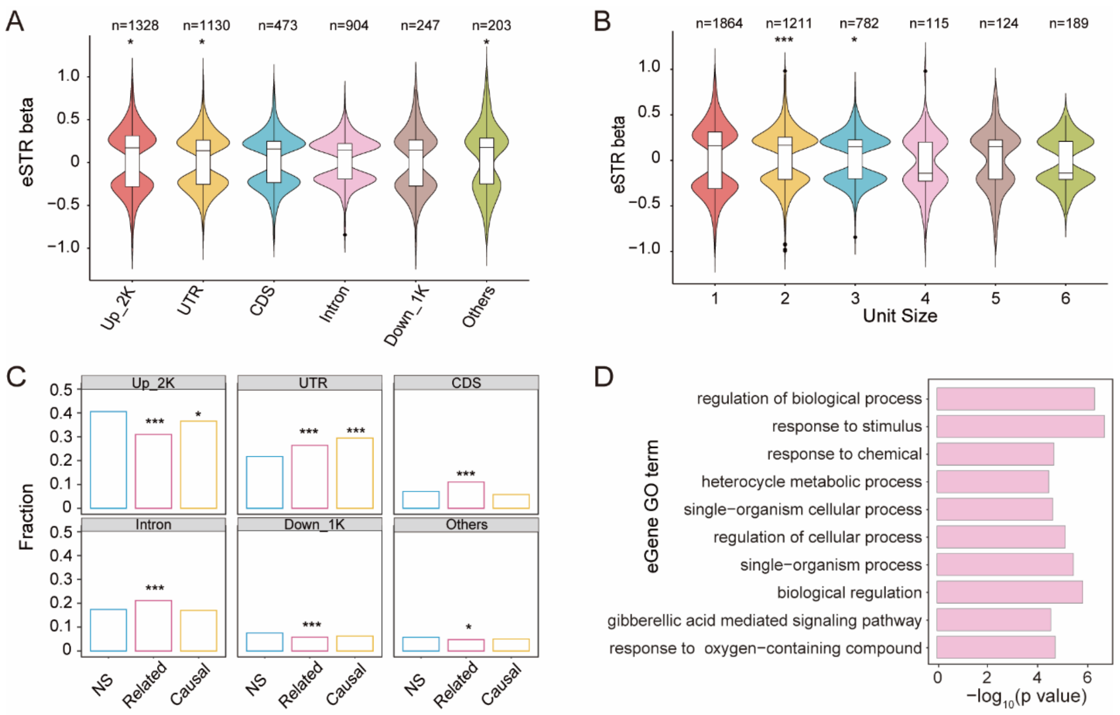


Figure S3. The effects of STRs on gene expression. (A) Effect size (beta value) of eSTRs located in different genomic regions. Beta value of the y-axis represents the correlation coefficient between STR length and expression variation, and a value greater than 0 indicates a positive correlation. Binomial test was used. *, $p < 0.05$; **, $p < 0.01$; ***, $p < 0.001$. (B) Effect size of eSTRs with different unit size. Binomial test was used. (C) Enrichment of STRs in different genomic regions of associated genes in eQTL analysis. NS, STRs that were not reported to be significantly related with any gene expression; related, STRs that were significantly associated with at least one gene expression variation but were not reported as causal; causal, STRs that were reported as causal for at least one gene expression variation. Fisher's exact test was used. (D) GO enrichment of genes with eSTRs (eGenes) by agriGO (Tian et al. 2017).

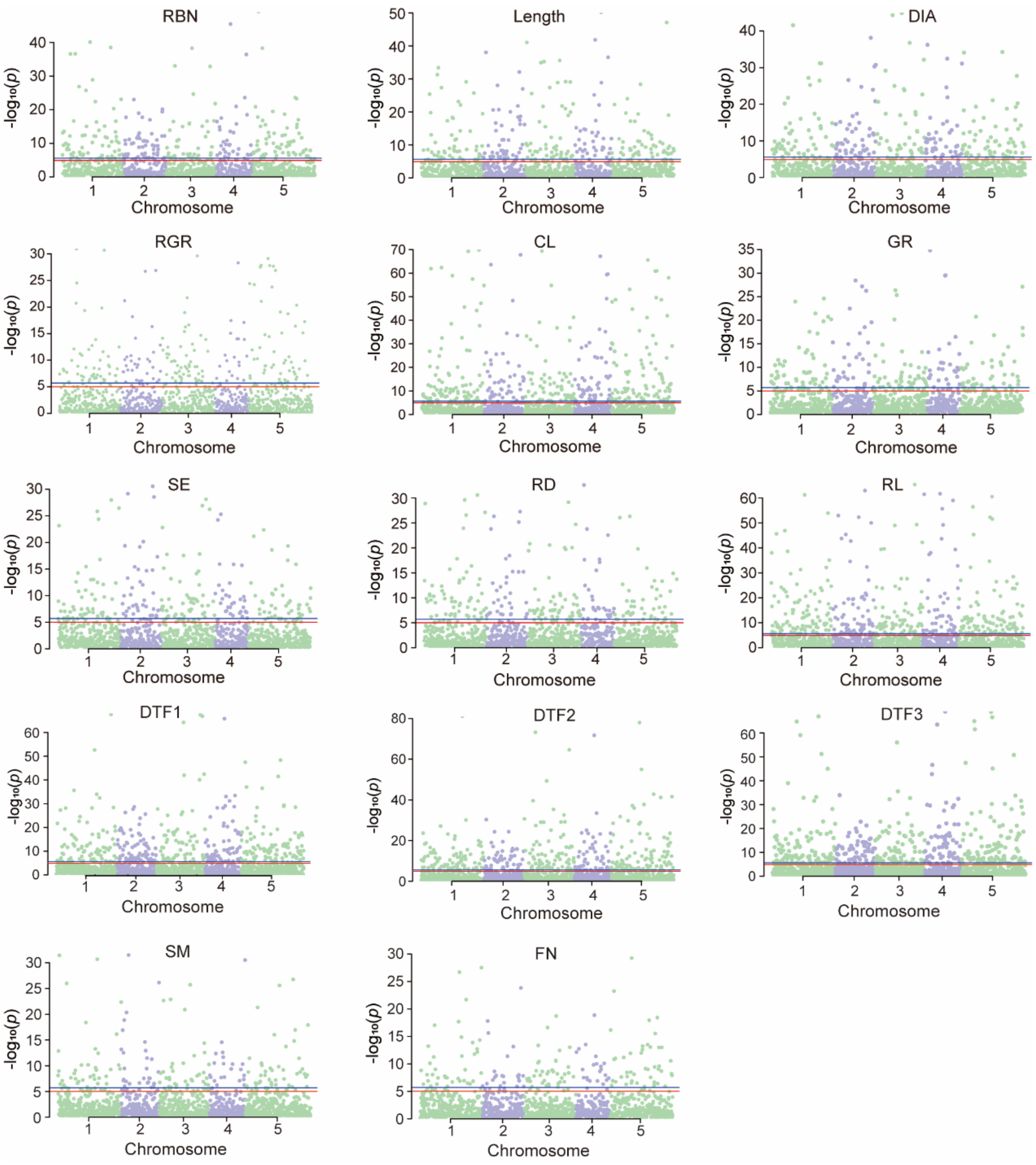


Figure S4. GWAS analysis using STR loci. Manhattan plots are shown for 14 phenotypes. The red horizontal line corresponds to the significance threshold (0.05/STR loci number) and blue horizontal line corresponds to the significance threshold (0.01/STR loci number).

Table S1. Summary of mutation rates of seven mutation accumulation lines (Weng et al. 2021), used in this study.

| Founder | CN1A18 | CN2A16 | RUD4 | RUD6 | SB4 | SB5 | Col-0 |
|---|---|---|---|---|---|---|---|
| No. of lines (L) | 56 | 51 | 50 | 50 | 53 | 56 | 107 |
| No. of generations (T) | 10 | 10 | 8 | 8 | 8 | 8 | 25 |
| Mutation rate (per locus per generation) | 1.93E-02 | 1.72E-02 | 6.05E-03 | 1.26E-02 | 5.46E-03 | 4.40E-03 | 5.78E-03 |

Table S2. Summary of mutation rates for STRs of different repeat unit.

| **STR repeat unit** | **μ** | **unit length** | **number** |
|---|---|---|---|
| A | 0.001775 | 1 | 120 |
| C | 0.002185 | 1 | 17 |
| AT | 0.002627 | 2 | 725 |
| AG | 0.000784 | 2 | 31 |
| AC | 0.000405 | 2 | 4 |
| ATC | 0.000424 | 3 | 6 |
| ATT | 0.000505 | 3 | 5 |
| CTT | 0.001785 | 3 | 11 |
| GTT | 0.000374 | 3 | 2 |
| AAAGC | 0.000748 | 5 | 1 |
| AAACAC | 0.000374 | 6 | 1 |
| AAGGTT | 0.000748 | 6 | 1 |
| AATGGC | 0.000374 | 6 | 1 |
| ACCACG | 0.000748 | 6 | 1 |

Table S3. Summary of the 10 assembled genomes used in this study.

| Data | Technology | References |
|---|---|---|
| Col-0 | Sanger sequencing | (The Arabidopsis Genome Initiative 2000) |
| 5-15 | PacBio | (Jiang et al. 2024) |
| KBS-Mac-74 | PacBio | (Michael et al. 2018) |
| An-1 | PacBio | (Jiao and Schneeberger 2020) |
| C24 | | |
| Cvi-0 | | |
| Eri-1 | | |
| Kyo | | |
| Ler | | |
| Sha | | |

Table S4. Summary of the published natural accessions used in this study.

| Source | Population | Accessions |
|---|---|---|
| 1001 (The 1001 Genomes Consortium 2016) | North America | 1612,1622,1651,1652,1676,1684,1756,1757,1793,1797,1820,1829,1834,1835,1851,1852,1872,1890,1925,2016,2017,2031,2053,2057,2108,2171,2191,2212,2239,2240,2278,2285,2286,506,544,546,628,630,6739,6740,6744,6749,6750,6805,6806,680,6814,681,685,687,6926,6927,7248,728,7350,7356,7358,7359,7383,7416,7475,7514,7515,7525,7529,7530,7568,7717,7757,7767,801,8037,8057,8171,8233,8246,8464,8483,853,854,867,868,870,9027,932 |
| 1001 | NW. China & C. Asia | 14312,14313,14314,14315,14318,14319,15560,18694,18696,6929,6931,6938,7183,7323,763,766,772,8424,9125,9128,9130,9131,9133,9134,9609,9611,9612,9616,9620,9621,9625,9626,9629,9630,9631,9635,9636,9639,9640,9641 |
| 1001 | Central Europe | 10020,10027,15591,15593,19949,19950,19951,403,410,424,428,430,5837,5874,5890,5893,5907,5921,5950,5984,5993,6008,6296,6390,6396,6424,6445,6903,6919,6951,6956,6957,6975,6976,6979,6984,7025,7067,7120,7177,7186,7203,7207,7236,7273,7347,7372,7396,7411,7413,7520,7521,8235,8236,8284,8285,8290,8311,8365,8386,8419,9644,9664,9665,9669,9670,9671,9672,9673,9677,9678,9679,9682,9683,9684,9685,9686,9689,9690,9692,9693,9694,9695,9696,9730,9731,9732,9733,9756,9768,9769,9770,9771,9772,9774,9775,9776,9777,9778,9779,9780,9781,9782,9784,9785,9786,9787,9788,9789,9791,9792,9793,9794,9795,9796,9797,9798,9799,9800,9801,9802,9803,9804,9805,9806,9807,9808,9812,9815 |
| 1001 | Germany | 5486,5748,5800,6180,6252,6268,6276,6915,6940,6945,6973,6997,7002,7008,7014,7036,7062,7102,7117,7119,7133,7143,7161,7165,7192,7199,7202,7208,7258,7276,7282,7287,7322,7419,7461,7516,8238,8312,9442,9476 |
| 1001 | Balkans | 7077,9067,9069,9070,9075,9078,9081,9084,9085,9089,9091,9095,9100,9102,9103,9104,9105,9106,9111,9113,9114,9115,9121,9613,9647,9649,9653,9656,9660,9698,9699,9700,9701,9703,9705,9708,9710,9711,9712,9713,9717,9720,9722,9726,9755 |
| 1001 | North Sweden | 1254,1257,1552,5856,5860,6009,6010,6011,6012,6013,6016,6017,6025,6046,6064,6069,6070,6071,6153,6154,6163,6166,6169,6172,6173,6174,6177,6184,6209,6210,6214,6216,6217,6218,6220,6221,6235,6238,6240,6244,6900,6901,6913,6917,6918,6968,6969,8227,8351,8376,9321,9323,9332,9363,9371,9386,9388,9427,9433 |
| 1001 | Admixed | 10022,10023,1313,1317,265,5720,5757,5768,5772,5784,5811,6039,6042,6086,6258,6434,6830,6907,6920,6932,6981,6992,7003,7028,7058,7061,7096,7111,7126,7127,7147,7163,7169,7181,7209,7213,7288,7305,7314,7316,7342,7353,7373,7384,7394,7404,8256,8334,8366,9058,9336,9343,9381,9382,9383,9394,9506,9508,9513,9523,9527,9529,9530,9536,9548,9551,9552,9558,9559,9561,9565,9576,9579,9581,9 |

| | | |
|---|---|---|
| | | 591,9595,9596,9663,9743,9754,9764,9783,9831,9835,9839,9849,9881,9890,9892,9894,9933 |
| 1001 | Relicts | 6911,9533,9542,9543,9545,9549,9550,9554,9555,9574,9583,9598,9600,9606,9832,9837,9871,9879,9887 |
| 1001 | South Sweden | 1002,1006,1061,1062,1063,1066,1070,1158,1166,5830,5831,5832,5836,5865,5867,6019,6020,6021,6022,6023,6024,6034,6036,6038,6040,6041,6073,6074,6076,6077,6085,6087,6088,6090,6091,6092,6094,6095,6096,6097,6098,6099,6100,6101,6102,6104,6105,6106,6107,6109,6111,6112,6113,6114,6115,6118,6119,6122,6123,6124,6125,6126,6128,6131,6132,6133,6134,6136,6137,6138,6140,6141,6142,6145,6148,6149,6150,6151,6188,6189,6191,6192,6193,6194,6195,6201,6202,6203,6242,6284,6413,6974,7013,7346,7354,8222,8230,8231,8234,8237,8240,8241,8242,8247,8249,8258,8259,8283,8306,8307,8326,8335,8369,8422,8426,8427,9057,9339,9352,9353,9369,9370,9380,9390,9391,9392,9399,9402,9404,9405,9407,9408,9409,9412,9413,9416,9421,9436,9451,9452,9453,9454,9455,9470,9471,9481,991,992,997 |
| 1001 | Spain | 6933,6961,6971,7081,7327,7328,8264,8357,9507,9509,9510,9511,9512,9514,9515,9518,9519,9520,9521,9522,9524,9525,9526,9531,9532,9534,9535,9537,9540,9541,9544,9546,9547,9553,9556,9557,9560,9562,9564,9567,9568,9573,9577,9582,9584,9586,9587,9588,9589,9593,9594,9597,9601,9602,9817,9819,9820,9821,9822,9825,9833,9834,9836,9840,9841,9843,9845,9846,9848,9850,9852,9855,9868,9870,9873,9874,9876,9878,9880,9888,9891,9895,9898,9900,9901,9902 |
| 1001 | Western Europe | 108,139,159,350,351,4807,4840,4857,4939,4958,5210,5249,5253,5276,5279,5349,5353,5395,5577,5644,5651,5717,5718,5726,5741,5776,5779,5798,6108,6904,6908,6923,6943,6944,6958,6959,6960,6966,6967,6986,7026,7064,7071,7092,7130,7217,7387,8214,8243,8244,8297,8337,88,9569,9571,9578,9585,9590,9599,9809,9826,9827,9847,9851,9853,9854,9875 |
| Africa | Relicts | Agl0,Agl1,Agl2,Agl3,Agl5,Agl9,Ait9,Ait14,Arb0,Arb2,Azr0,Azr5,Azr7,Azr11,Azr13,Azr16,Bab0,Bab3,Bba0,Bba2,Bbe0,Elh2,Elh10,Elh15,Elh20,Elh23,Elh27,Elh33,Elh39,Elh46,Elk1,Elk3,Elk20,Elk28,IFr0,Ifr3,Ifr4,Ifr6,Ket10,Ket12,Khe0,Khe32,Meh0,Meh4,Meh7,Oua0,Set0,Set6,Tah0,Tah4,Tanz-1,Taz0,Taz11,Taz16,Taz18,Til2,Tiz0,Tiz7,Zin4,Zin9 |
| China (Zou et al. 2017; Jiang et al. 2024) | NW. China & C. Asia | 39-16,40-2,41-10,41-11,42-3,42-8,43-12,43-7,45-20,45-23,46-28,46-31,48-1,50-3,50-5,51-1,51-4,51-8,52-10,52-13,53-18,53-19,54-5,54-6,55-12,55-13,55-22, 39-6,39-7,41-7,42-4,43-1,45-25,46-27,50-4,53-21,54-4,55-7 |
| China | Relicts | 36-31,87 |
| China | Yangtze | 10-1,10-3,10-5,11-4,12-8,13-13,1-32,13-5,14-62,15-11,17-2,17-5,18-4,18-7,1-8,19-33,20-2,20-36,23-19,23-34,24-10,25-16,2-5,26-5,27-8,27-9,28-6,29-8,30-1,30-6,31-26,32-28,3-2,33-46,33-4,34-16,3-4,35-10,35-1,37-10,38-2,38-4,4-15,5-15,57-1,58-7,59-12,60-1,61-1,62-1,6-28,63-1,64-5,65-1,66-3,67-1,68-1,69-7,6-9,70-1,71-5,72-2,7-2,73- |

| | | |
|---|---|---|
| | | 5,74-14,75-6,76-3,76-4,76-5,77-2,77-3,78-4,78-5,79-2,79-4,8-17,82-1,82-3,83-2,83-5,84-3,84-4,85-3,85-5,86,8-6,9-5, 11-1,11-5,12-11,12-3,13-10,14-5,16-62,17-11,18-5,19-61,19-66,20-21,21-42,21-54,21-64,2-16,2-1,22-10,22-20,23-40,24-14,24-20,25-30,25-34,26-1,26-3,27-5,28-10,28-17,29-50,29-6,30-7,3-1,32-15,32-27,33-53,34-11,34-2,35-3,37-3,37-8,38-3,4-17,4-7,5-23,57-6,57-7,58-2,58-5,59-2,59-4,60-2,60-7,61-2,61-5,62-2,62-3,63-2,63-5,64-2,64-3,65-2,65-4,66-5,66-7,68-3,68-5,6-8,69-1,69-5,70-2,70-5,71-1,71-4,7-15,7-1,72-4,72-6,73-9,74-10,74-8,75-3,75-5,77-5,78-1,79-6,80-1,80-2,81-10,81-2,81-3,82-6,83-3,84-1,85-1,88-1,88-2,88-3,8-8,89-1,89-2,89-5,90-1,90-2,90-6,9-10,91-1,9-15,92-2,92-4,92-6,93-2,93-5,93-6,94-1,94-2,95-3 |

Note: NW. China & C. Asia, northwestern China and central Asia population; Yangtze, Yangtze River basin population.

Table S5. Summary of the 55 natural accessions resequenced in this study.

| Population | Accession | Region | Latitude | Longitude |
|---|---|---|---|---|
| Yangtze | 1-13 | Ningguo, Anhui | 30.68 | 118.97 |
| Yangtze | 15-14 | Chun'an, Zhejiang | 29.8 | 118.85 |
| Yangtze | 22-1 | Pan'an, Zhejiang | 28.96 | 120.37 |
| Yangtze | 73-6 | Xingzi, Jiangxi | 29.29 | 115.98 |
| Yangtze | 91-2 | Jinmen, Hubei | 30.55 | 113.77 |
| Yangtze | 96-1 | Shangrao, Jiangxi | 28.24 | 118.05 |
| Yangtze | 96-5 | Shangrao, Jiangxi | 28.24 | 118.05 |
| Yangtze | 97-3 | Shangrao, Jiangxi | 28.24 | 118.05 |
| Yangtze | 97-5 | Shangrao, Jiangxi | 28.24 | 118.05 |
| Yangtze | 98-1 | Jinhua, Zhejiang | 29.12 | 119.73 |
| Yangtze | 99-2 | Zhuji, Zhejiang | 29.72 | 120.19 |
| Yangtze | 102-1 | Jingxian, Anhui | 30.69 | 118.38 |
| Yangtze | 102-3 | Jingxian, Anhui | 30.69 | 118.38 |
| Yangtze | 102-5 | Jingxian, Anhui | 30.69 | 118.38 |
| Yangtze | 103-6 | Jingxian, Anhui | 30.67 | 118.38 |
| Yangtze | 104-1 | Jinzhai, Anhui | 31.63 | 116 |
| Yangtze | 105 | Jinzhai, Anhui | 31.62 | 116.01 |
| Yangtze | 106-1 | Jinzhai, Anhui | 31.62 | 116.01 |
| Yangtze | 109-1 | Xuancheng, Anhui | 30.96 | 118.77 |
| Yangtze | 109-5 | Xuancheng, Anhui | 30.96 | 118.77 |
| Yangtze | 110-2 | Huoshan, Anhui | 31.46 | 116.37 |
| Yangtze | 112-2 | Yuexi, Anhui | 30.92 | 116.35 |
| Yangtze | 113 | Yuexi, Anhui | 30.92 | 116.35 |
| Yangtze | 114-5 | Yuexi, Anhui | 30.58 | 116.59 |
| Yangtze | 115-1 | Qianshan, Anhui | 30.58 | 116.6 |
| Yangtze | 116-3 | Qianshan, Anhui | 30.57 | 116.6 |
| Yangtze | 117 | Qianshan, Anhui | 30.7 | 116.57 |
| Yangtze | 118-1 | Qianshan, Anhui | 30.57 | 116.61 |
| Yangtze | 118-4 | Qianshan, Anhui | 30.57 | 116.61 |
| Yangtze | 119-1 | Qianshan, Anhui | 30.71 | 116.57 |
| Yangtze | 119-2 | Qianshan, Anhui | 30.71 | 116.57 |
| Yangtze | 119-3 | Qianshan, Anhui | 30.71 | 116.57 |
| Yangtze | 120 | Qianshan, Anhui | 30.71 | 116.57 |
| Yangtze | 121-3 | Shitai, Anhui | 30.23 | 117.46 |
| Yangtze | 122-5 | Shitai, Anhui | 30.2 | 117.5 |
| Yangtze | 124-2 | Huangshanqu,Anhui | 30.2 | 118.15 |
| Yangtze | 124-4 | Huangshanqu,Anhui | 30.2 | 118.15 |
| Yangtze | 125-2 | Huangshanqu,Anhui | 30.23 | 118.11 |
| Yangtze | 125-5 | Huangshanqu,Anhui | 30.23 | 118.11 |
| Yangtze | 126-3 | Huangshanqu,Anhui | 30.23 | 118.14 |

| | | | | |
|---|---|---|---|---|
| Yangtze | 127-1 | Huangshanqu,Anhui | 30.3 | 118.14 |
| Yangtze | 127-2 | Huangshanqu,Anhui | 30.3 | 118.14 |
| Yangtze | 128-4 | Huangshanqu,Anhui | 30.29 | 118.1 |
| Yangtze | 131-1 | Jingde, Anhui | 30.33 | 118.54 |
| Yangtze | 131-3 | Jingde, Anhui | 30.33 | 118.54 |
| Yangtze | 131-5 | Jingde, Anhui | 30.33 | 118.54 |
| Yangtze | 132-1 | Jingde, Anhui | 30.32 | 118.53 |
| Yangtze | 132-5 | Jingde, Anhui | 30.32 | 118.53 |
| Yangtze | 133-3 | Jingde, Anhui | 30.33 | 118.53 |
| Yangtze | 133-5 | Jingde, Anhui | 30.33 | 118.53 |
| Yangtze | 134-3 | Jingde, Anhui | 30.33 | 118.53 |
| Yangtze | 134-4 | Jingde, Anhui | 30.33 | 118.53 |
| Yangtze | 137-1 | Ningguo, Anhui | 30.62 | 119 |
| Yangtze | 137-2 | Ningguo, Anhui | 30.62 | 119 |
| Yangtze | 138-1 | Chaohu, Anhui | 31.66 | 117.72 |

Note: Yangtze, Yangtze River basin population.

Table S6. Summary of STR identification accuracy using short-read data.

| Accession | 5-15 | Cvi-0 | Kyo | Ler |
|---|---|---|---|---|
| No. of STRs identified in short reads | 30814 | 24249 | 13456 | 21234 |
| No. of STRs confirmed in assembled data | 30143 | 23317 | 13009 | 20441 |
| Accuracy ratio (%) | 97.82% | 96.16% | 96.68% | 96.27% |

Table S7. Summary of the published phenotypes used in this study (Togninalli et al. 2020).

| Phenotypes | Phenotype Description | Growth conditions |
|---|---|---|
| FT10 | Flowering time (FT) | 10℃, 16 hours daylight |
| FT16 | Flowering time (FT) | 16°C, 16 hours daylight |
| DTF1 | Days Until Emergence of Visible Flowering Buds in the Center of the Rosette from Time of Sowing | Plants were grown in controlled growth chambers with the following settings: 16 h light/8 h darkness, 16°C constant temperature, 65% humidity |
| DTF2 | Days Until the Inflorescence Stem Elongated to 1cm | |
| DTF3 | Days Until First Open Flower | |
| CL | Cauline Leaf Number | |
| RL | Rosette Leaf Number | |
| Length | Length of Main Flowering Stem | |
| RBN | Rosette Branch Number | |
| DIA | Diameter of Rosette (End point after flowering) | |
| SM | Stomata size was measured using high-throughput microscopy from 7-week-old | Plants were grown for 7 weeks in growth chambers (one per block) under the following conditions: 16 hr light; 95μmol $s^{-1}mm^{-2}$ light intensity; and 20°Cday-and 18°C night temperature |
| GR | Average growth rate during the whole life cycle (final dry mass / lifespan, mg d-1) | Growth chamber set at 16°C 12/12 h photoperiod, under semi-hydroponic conditions |
| RD | Rosette dry mass (mg) | Growth chamber set at 16°C 12/12 h photoperiod, under semi-hydroponic conditions. |
| SE | Measured as first derivative of the scaling relationship between growth rate and plant dry mass | |
| RGR | Relative growth rate (mg d-1 g-1), half final plant size | |
| FN | Total number of fruits per plant at the end of reproduction (fruit ripening) | |
